# Functional and Comparative Genomics of Niche-Specific Adapted Actinomycetes *Kocuria rhizophila* Strain D2 Isolated from Healthy Human Gut

**DOI:** 10.1101/400242

**Authors:** Vikas C. Ghattargi, Yogesh S. Nimonkar, Kamala Sape, Om Prakash, Mangesh V. Suryavanshi, Yogesh S. Shouche, Bharati S. Meti, Shrikant P. Pawar

## Abstract

Incidences of infection and occurrence of *Kocuria rhizophila* in human gut are prominent but certainly no reports on the species ability to withstand human gastrointestinal dynamics. *Kocuria rhizophila* strain D2 isolated from healthy human gut was comprehensively characterized. The functional analysis revealed the ability to produce various gastric enzymes and sensitive to major clinical antibiotics. It also exhibited tolerance to acidic pH and bile salts. Strain D2 displayed bile-salt hydrolytic (BSH) activity, strong cell surface traits such as hydrophobicity, auto-aggregation capacity and adherence to human HT-29 cell line. Prominently, it showed no hemolytic activity and was susceptible to the human serum. Exploration of the genome led to the discovery of the genes for the above said properties and has ability to produce various essential amino acids and vitamins. Further, comparative genomics have identified core, accessory and unique genetic features. The core genome has given insights into the phylogeny while the accessory and unique genes has led to the identification of niche specific genes. Bacteriophage, virulence factors and biofilm formation genes were absent with this species. Housing CRISPR and antibiotic resistance gene was strain specific. The integrated approach of functional, genomic and comparative analysis denotes the niche specific adaption to gut dynamics of strain D2. Moreover the study has comprehensively characterized genome sequence of each strain to know the genetic difference and intern recognize the effects of on phenotype and functionality complexity. The evolutionary relationship among strains along and adaptation strategies has been included in this study.

**Significance:** Reports of Kocuria rhizophila isolation from various sources have been reported but the few disease outbreaks in humans and fishes have been prominent, but no supportive evidence about the survival ability of Kocuria spp. within human GIT. Here, we report the gut adaption potential of K. rhizophila strain D2 by functional and genomic analysis. Further; comparative genomics reveals this adaption to be strain specific (Gluten degradation). Genetic difference, evolutionary relationship and adaptation strategies have been including in this study.

## INTRODUCTION

The genus Kocuria (formerly Micrococcus) was named after a Slovakian microbiologist: Miroslav Kocur, and belongs to the class *Actinobacteria* (1). It is Gram-positive cocci, catalase positive, non-hemolytic, non-endospore-forming, non-motile and can grow at different oxygen levels (aerobic, facultative and anaerobic) (1, 2). Presence of galactosamine and glucosamine (amino sugars) as the main component of the cell wall, differ them from other members of the class *Actinobacteria* (2). This genus is normally inhabitants of dust, soil, water and food and in humans colonizes skin, mucosa, oropharynx and gastrointestinal tract (GIT) (3–5).

*K. rhizophila* was isolated from the rhizoplane of the narrow-leaved cattail (*Typha angustifolia*). in 1999 (6). The strain ATCC 9341 was reclassified as *K. rhizophila* from *Micrococcus luteus* (7). It has been isolated from viz. cheese (8), chicken meat (9) and also healthy human GIT (1) across the globe, thus suggesting its wide adaption potential. Moreover, it has been important in industrial applications for antimicrobial susceptibility testing as standard quality control strain (1–3).

Currently, *K. rhizophila* is gaining importance as emerging pathogen in immune-compromised and metabolically disordered individuals (10–14). In particular, their affinity to plastic materials and devices such as a catheter, causing chronic recurrent bacteremia and thus causing mortality (10–14). Therefore, one should not underestimate the significance of such microorganisms when isolated from clinical samples and particularly from Gastrointestinal tract (GIT), blood and medical implant surfaces. Currently, there are two complete genomes (*K. rhizophila* DC2201 and FDAARGOS_302) and five draft genome (P7-4, TPW45, 14ASP, RF and G2) sequences of *K. rhizophila* available publicly at NCBI (National Center for Biotechnology Information). Based on genome sequence it displays a wide range of activity viz. tolerance to various organic compounds and sturdy amino-acid and carbohydrate metabolism.

Recent techniques of 16S rRNA amplicon and metagenomics sequencing have vividly expanded the known diversity of the human gut microbiome (15–18) but the first approach used to study the gut microbiota employed microbial culture (1). Recent studies with culturomics approach have provided actual insights into the type of species present in the human gut (18–20). Using above-said method (culturomics), we could isolate more than 120 different strains from 18 different genera were isolated; using 35 culture media and different growth conditions.

Here, we present the work carried out on *K. rhizophila* strain D2 isolated from the healthy human gut as there is no supportive evidence about the survival ability of *Kocuria spp.* within human GIT. Thus we found the importance to study strain D2 for its adaption, pathogenicity, commensal or beneficial nature. The work described here utilizes *in-vitro*, genomes and comparative genomics approach to identify the potential of *K. rhizophila* strain D2.

## RESULTS

### Identification

The 16S rRNA gene of strain D2 showed 99.92% similarity to *Kocuria rhizophila* type strain DSM 11926. The phylogenetic tree was constructed using the Neighbour-Joining method with closely related taxa (Fig 1).

**Fig 1.**
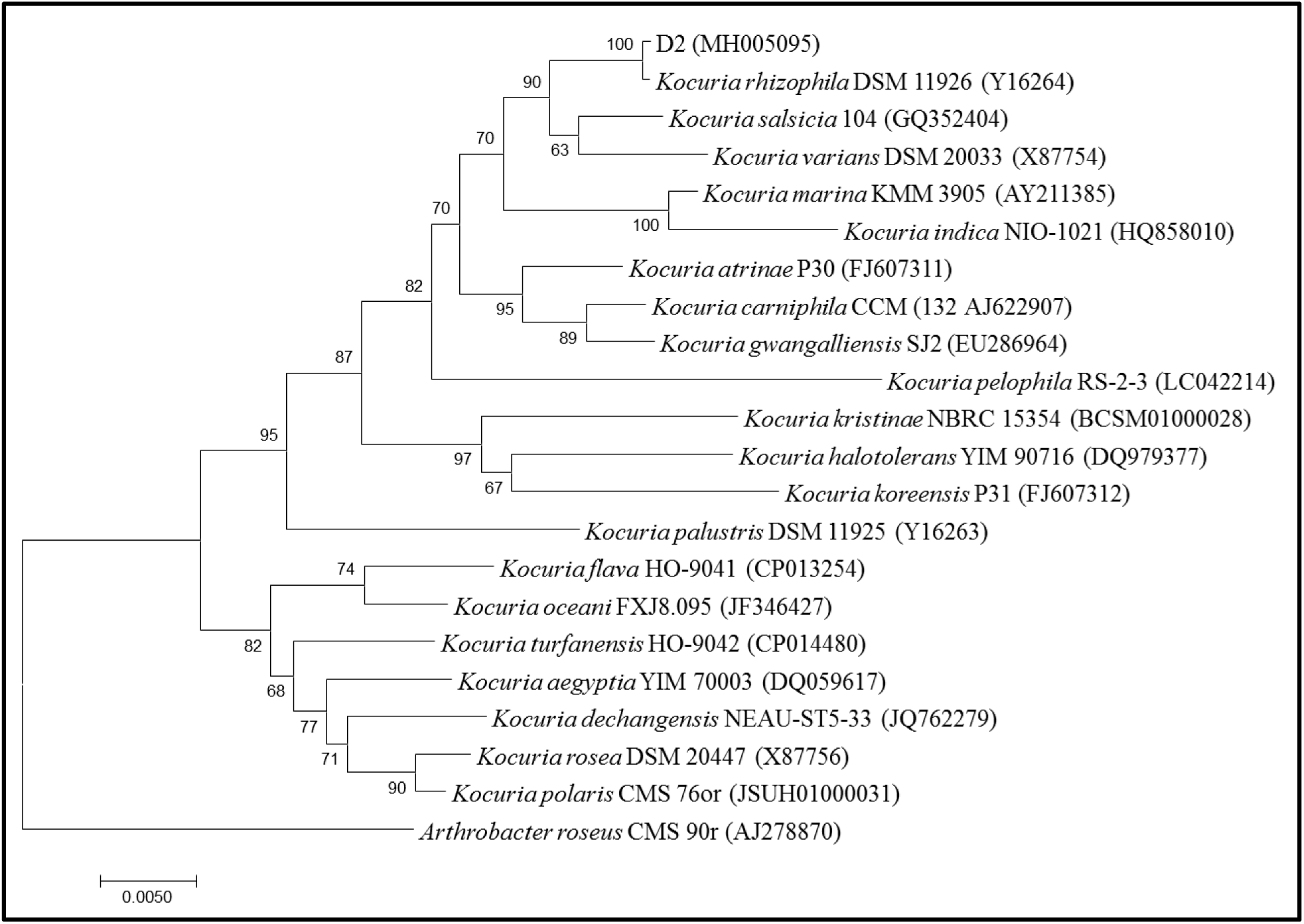
Phylogenetic relationship of strain D2 with closely related taxa based on 16S rRNA gene sequences (16S rRNA gene sequence of *Arthrobacter roseus* was used as out-group). The phylogenetic trees were constructed using the Neighbour-Joining method. The evolutionary distances were computed using the Kimura 2-parameter method and are in the units of the number of base substitutions per site. The rate variation among sites was modelled with a gamma distribution (shape parameter = 1)

### Characterization of *K. rhizophila* D2

#### Exoenzymes, Carbohydrate utilization, Antibiogram and Plasmid determination

The strain D2 showed various gastric enzymes viz. lipase, urease, phosphatase, protease, catalase and amylase. The activity was also found for nitrate reductase activities (Table 1). The strain was also able to utilize most of the sugars viz. inulin, lactose, sucrose, fructose, maltose, galactose, dextrose, raffinose, trehalose and melibiose (Table 2). The strain had intermediate resistance to levofloxacin, ciprofloxacin and gentamicin while sensitive to other 21 of the 24 tested antibiotics (Table 3). The plasmid was not associated with the strain.

**Table 1:**
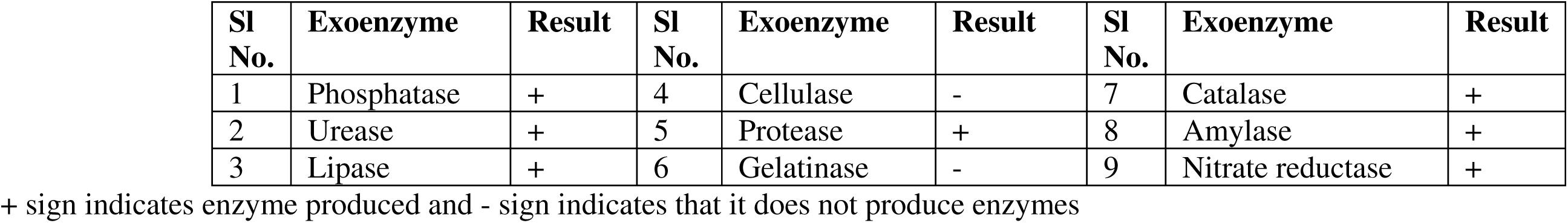
Exoenzymes produced by *Kocuria rhizophila* D2

| Sl No. | Exoenzyme | Result | Sl No. | Exoenzyme | Result | Sl No. | Exoenzyme | Result |
| --- | --- | --- | --- | --- | --- | --- | --- | --- |
| 1 | Phosphatase | + | 4 | Cellulase | - | 7 | Catalase | + |
| 2 | Urease | + | 5 | Protease | + | 8 | Amylase | + |
| 3 | Lipase | + | 6 | Gelatinase | - | 9 | Nitrate reductase | + |

+ sign indicates enzyme produced and - sign indicates that it does not produce enzymes

**Table 2:** Various carbohydrates utilised by test strain *Kocuria rhizophila* D2

| Sl. No. | Carbohydrates | Strain D2 | Sl. No. | Carbohydrates | Strain D2 |
| --- | --- | --- | --- | --- | --- |
| 1 | Lactose | Y | 18 | Inositol | Y |
| 2 | Xylose | N | 19 | Sorbitol | Y |
| 3 | Maltose | Y | 20 | Mannitol | Y |
| 4 | Fructose | Y | 21 | Adonitol | N |
| 5 | Dextrose | Y | 22 | Arabitol | Y |
| 6 | Galactose | Y | 23 | Erythritol | N |
| 7 | Raffinose | Y | 24 | Rhamnose | Y |
| 8 | Trehalose | Y | 25 | Cellobiose | Y |
| 9 | Melibiose | Y | 26 | Melezitose | N |
| 10 | Sucrose | Y | 27 | alpha-Methyl-D-Mannoside | N |
| 11 | L-Arabinose | Y | 28 | Xylitol | N |
| 12 | Mannose | N | 29 | ONPG | N |
| 13 | Inulin | Y | 30 | Esculin | Y |
| 14 | Sodium gluconate | N | 31 | D-Arabinose | Y |
| 15 | Glycerol | N | 32 | Citrate | Y |
| 16 | Salicin | N | 33 | Malonate | Y |
| 17 | Dulcitol | N | 34 | Sorbose | Y |

5 Y = Yes indicates its ability to utilise carbohydrate and N = No indicates could not utilise carbohydrate.

**Table 3:** Showing Antibiogram results for the strain under study. S: Sensitive, I: Intermediate and R: Resistance to tested antibiotics

| Sl. No | Antibiotic tested | Conc. tested | Strain D2 | Sl. No | Antibiotic tested | Conc. tested | Strain D2 |
| --- | --- | --- | --- | --- | --- | --- | --- |
| 1 | Cefpodoxime | 10µg | S | 13 | Amikacin | 30µg | S |
| 2 | Chloramphenicol | 30µg | S | 14 | CoTrimoxazole | 25µg | S |
| 3 | Vancomycin | 30µg | S | 15 | Colistin | 10µg | S |
| 4 | Streptomycin | 10µg | S | 16 | Augmentin | 30µg | S |
| 5 | Rifampicin | 5µg | S | 17 | Netillin | 30µg | S |
| 6 | Levofloxacin | 5µg | I | 18 | Norfloxacin | 10µg | S |
| 7 | Ceftriaxone | 30µg | S | 19 | Ceftriaxone | 10µg | S |
| 8 | Clindamycin | 2µg | S | 20 | Ciprofloxacin | 5µg | I |
| 9 | Augmentin | 30µg | S | 21 | Cefotaxime | 30µg | S |
| 10 | Amikacin | 30µg | S | 22 | Gentamicin | 10µg | I |
| 11 | Cefixime | 5µg | S | 23 | Furazolidone | 50µg | I |
| 12 | Tetracycline | 30µg | S | 24 | Amoxycillin | 10µg | S |

#### Bile, Acid and hydrogen peroxide tolerance

Strain D2 was able to grow in the various bile concentration from 0.1- 1 % (w/v), of these it showed 60% survivability at 0.4% bile salt after 24hrs. For the acid tolerance assay strain D2 showed 58.8% survivability at pH 2.5 for 3h. The time depicts the time taken by the food in the stomach. Further, strain D2 was able to tolerate hydrogen peroxide for 4.5 hours.

#### Adhesion, Exopolysaccharide production and Bile salt hydrolytic (BSH) activity

Auto-aggregation capacity of strain D2 was 29%, while the cell surface hydrophobicity to hydrocarbons viz. toluene, xylene and hexane was 31%, 28% and 22% respectively. The percent cell adhesion assay for human HT-29 cell line was found to be 30%. An additional microscopic observation revealed low adhesion across the quadrants (Fig 2). The strain D2 was able to hydrolyze bile, a zone of clearance (9 mm) was seen when grown on medium with bile salts and this ability was further confirmed by the Ninhydrin method [4 ±0.2 mg/ cell pellet (mg)]. The strain D2 could not produce exopolysaccharide.

**Fig 2.**
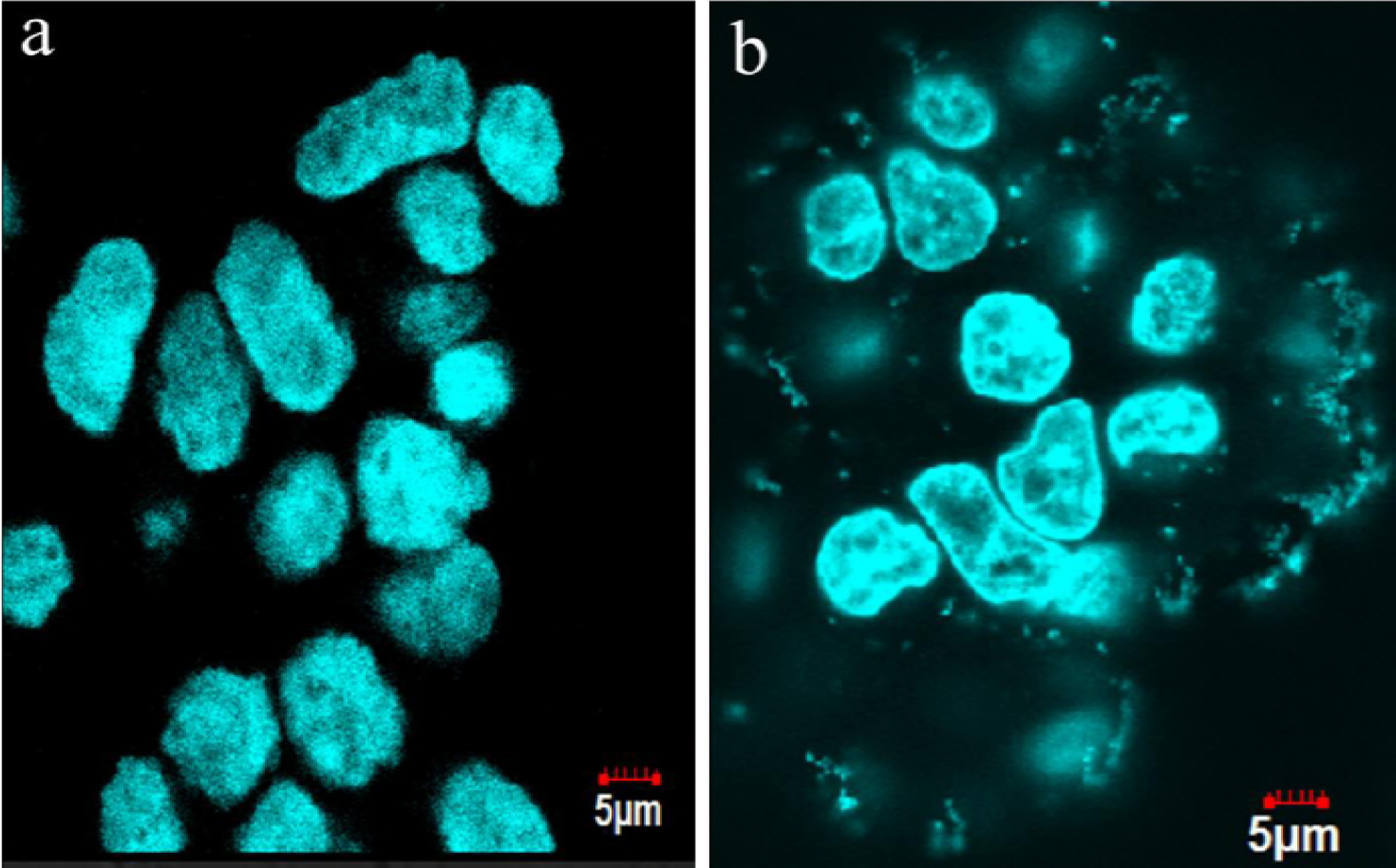
Results for cell adhesion assay performed on Human HT-29 Cell line. Images taken under fluorescence microscope 60X oil immersion where panel (a) is negative control without any bacterial cell added; panel (b) is for the strain D2

#### Resistance to Simulated GI Conditions and Gluten Degradation

The survival rate of strain D2 after 3 h exposure to simulated gastric juice was 78 × 10^8^ cells, which has reduced from 90 × 10^8^, the zero hour reading. Further, these cells were washed in PBS and added to the simulated intestinal juice. Only 59 × 10^8^ was alive after three hours. It could also utilize gluten as a sole source of nitrogen, and thus form a clear zone (11±0.1mm) around the colony grown.

#### Pathogenicity testing

Pathogenicity testing was included in the investigation. Hemolytic activity, serum resistance and biofilm formation have generally been used. The strain D2 was susceptible to human serum with 0.04% survival, not a biofilm producer and exhibited alpha hemolytic activity.

### Genome Features of *K. rhizophila* D2

More than 4.9 million good quality paired-end reads were obtained, with an approximate 110x sequencing coverage. The nearly complete genome of *Kocuria rhizophila* D2 consisted of 2,313,294 bp (2.3 Mb) with an average G+C content of 71.0 %. Genome consisted of 2,253 genes and 2,218 ORFs were identified. Within the genome, 46 structural tRNAs, and 3 rRNAs could be predicted.

### Genome-based metabolic capabilities of *K. rhizophila* D2

We screened the genome sequence of the strain D2 for various metabolic capabilities occurring within, helping to understand the cellular processes.

#### *Amino Acid Synthesis* and *Proteolytic System*

Analysis of genome reports the ability to synthesize amino acids such as serine, cysteine, and aspartate. From these amino acids, seven other could be generated. Complete pathways for essential amino acids such as valine and leucine could be constructed from the genome sequence. Only D-alanyl-D-alanine carboxypeptidase (EC 3.4.16.4) and three general peptidase genes could be identified in strain D2. Membrane proteins belonging to (Zinc) metalloendopeptidases was detected. Two copies of *Clp* protease (serine peptidases) were identified. The presence of aminopeptidases (E.C. 3.4.11.1 and E.C 3.4.11.2) and cytosol aminopeptidase *Pep*A were implied in gluten metabolism, found during the gluten utilization experiment.

#### Carbohydrate Metabolism

The genome of strain D2 encodes a large diversity of genes related to carbohydrate metabolism. Annotation from PATRIC and PGAP has shown the genes present in pyruvate metabolism I & II, glycolysis and gluconeogenesis, TCA cycle, pentose phosphate pathway and serine-glyoxylate cycle. The genes present in strain D2 helps to utilize various sugars viz. lactose, maltose, inositol, sorbitol, mannitol, fructose, adonitol, dextrose, arabitol, galactose, raffinose, rhamnose, trehalose, cellobiose, melibiose, sucrose, L-arabinose, esculin, d-arabinose, citrate, malonate and sorbose. The genomic analysis further countersigns the presence of these genes and their transporters within the genome.

#### Allied Metabolism

We could identify the entire components of thioredoxin system along with FMN reductase (EC 1.5.1.29) responsible for the sulfur assimilation into the cell. Enzymes exopolyphosphatase (EC 3.6.1.11) and polyphosphate kinase 2 (EC 2.7.4.1) catalyzes the hydrolysis of inorganic polyphosphate and formation of polyphosphate from ATP respectively. The genome also had seven genes responsible for ammonia assimilation into its cells, which chiefly includes glutamine synthesis genes. The genome also has pathways for the production of vitamins: menaquinone (vitamin K2), phylloquinone (vitamin K1), thiamine (B1) riboflavin (B2) and folate (B9). Also, complete pathways for carotenoids and primary bile salts could be identified.

#### Stress Response

The genome of *Kocuria rhizophila* D2 encodes various stress-related proteins, involving various proteases involved in the stress response. Highly conserved class I heat-shock genes (GroES and GroES operons) and F_1_F_0_-ATPase system for maintaining the integrity of cellular proteins under stress conditions were identified. The genome harbours three copies of universal stress proteins, non-specific DNA-binding protein and genes related to oxidative and osmotic stress were found.

### Comparative genomics

#### General genomic features

The average genome sizes of *K. rhizophila* strains were approximately of 2.74Mb and average GC content is 70.81 %. Comparison between GC content, genome sizes, number of genes and coding DNA sequence (CDS), we could not obtain any significant differences (P ≤ 0.05, Kruskal–Wallis statistical test). The average number of annotated protein-encoding genes is 2345. Analysis of RAST suggests the abundance of amino acids and its derivatives. Table 4 provided the general genome features for the strains.

**Table 4:**
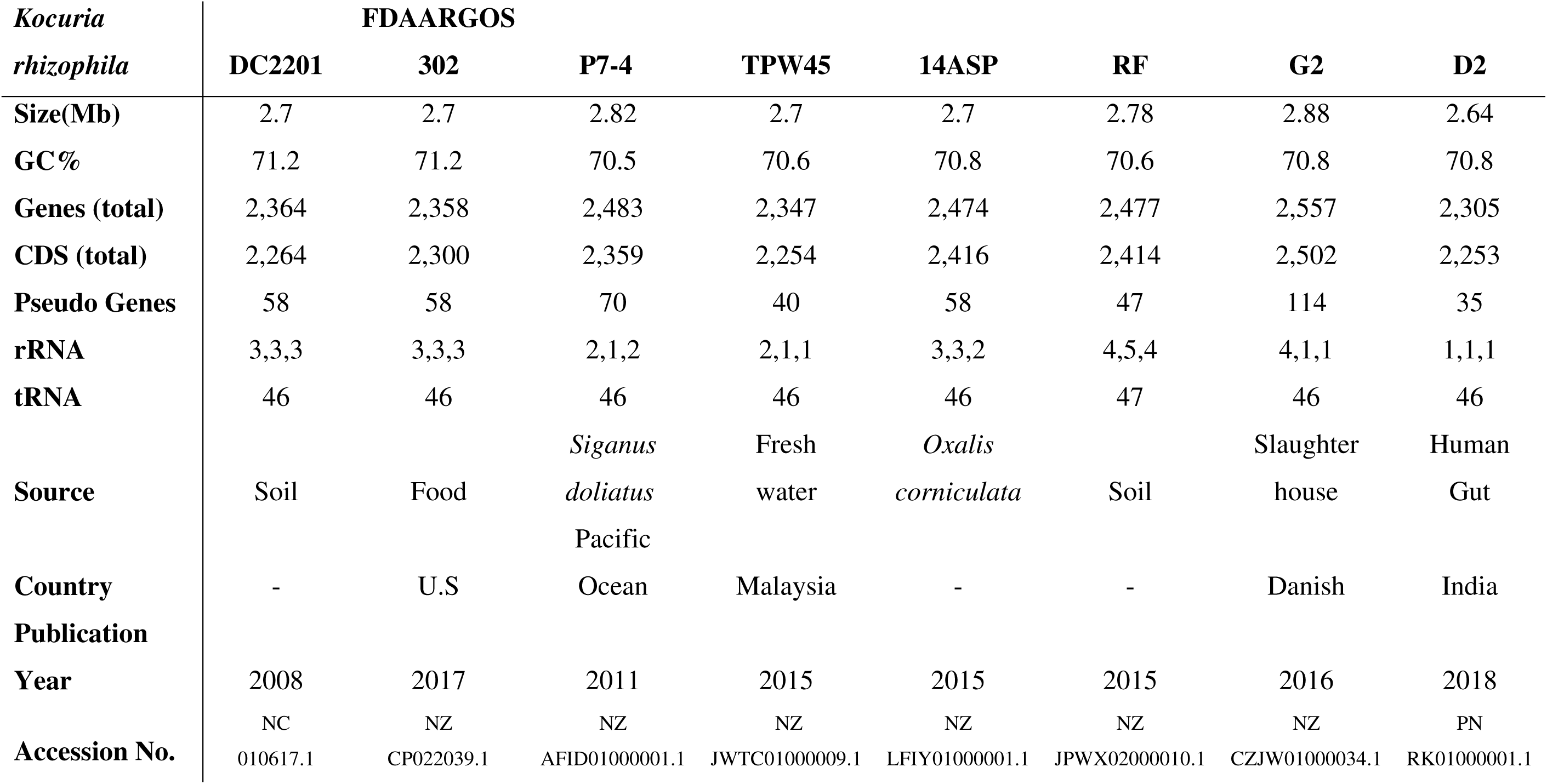
General genome features of various *Kocuria rhizophila* strains.

#### Comparisons of D2 with other K. rhizophila strains

The availability of various *K. rhizophila* genomes has helped to define a core, accessory, unique genome features. The comparison of strain D2 with other strains revealed 888 (40.11%) core genes, 1243 (56.14 %) accessory and 83(3.74%) unique genes. The core and pan graph for a number of shared gene families to the number of strains is plotted as shown in Fig 3. The size of the core-genome gradually stabilized while the pan-genome size grew continuously, by the addition of other strains indicating an open pan-genome.

**Fig 3.**
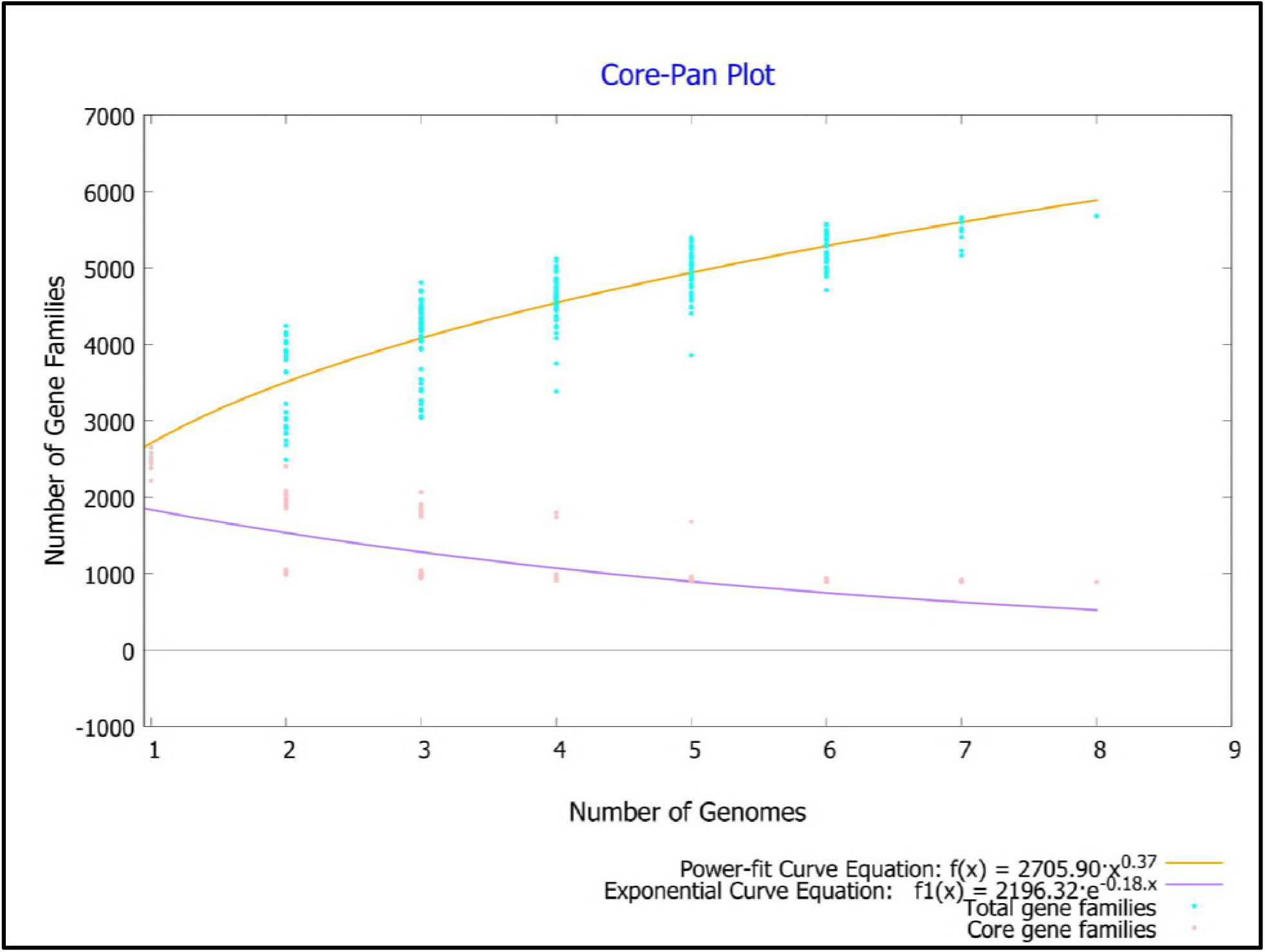
Core and pan genome for *K. rhizophila* strains. The number of shared genes is plotted as the function of number of strains (n) added sequentially. 888 copy genes are shared by all 8 genomes. The orange line represents the least-squares fit to the power law function f(x)=a.x^b where a= 2705.9, b= 0.373747. The red line represents the least-squares fit to the exponential decay function f1(x)=c.e^(d.x) where c= 2196.32, d= −0.179618.

#### Pan-Genome Analysis

The analysis of pan, core and accessory genome revealed the presence of 888 core, 10882 accessory genes. The number of strain-specific genes for strain 14ASP is 453, strain D2 are 83, DC2201 are 21, FDAARGOS_302 are 35, G2 are 514, P7-4 are 276, RF are 160 and for strain TPW45 are 203 (Fig 4). orthoMCL analysis of **core genes** leads to the identification of copy number within the genomes. 550 genes were present in a single copy and 338 genes were present in multiple copies within eight strains. Functional analysis of the core genes by Cluster of Orthologous Genes (COG) showed the distributed in a varied range of functional categories. These included genes related to cell growth, DNA replication, transcription, translation, and also general transporters. The analysis also revealed presence of carbohydrate, amino acid metabolism, stress response and secondary metabolism. Categories of representing growth, replication, transcription, translation, transporters comprised of 45.12% of the core genes. The core (888) genes were used to construct a phylogenetic tree for all the strains under study, where *K. flava* was used as an out-group. Phylogenetic reconstruction by using (ML) Maximum likelihood method separated the study 8 strains in 2 clusters with bootstrap more than 70 (Fig 5) and the same observation was made when repeated with the pan-genome (data not shown). Clustering based on the source of isolation could not be observed.

**Fig 4.**
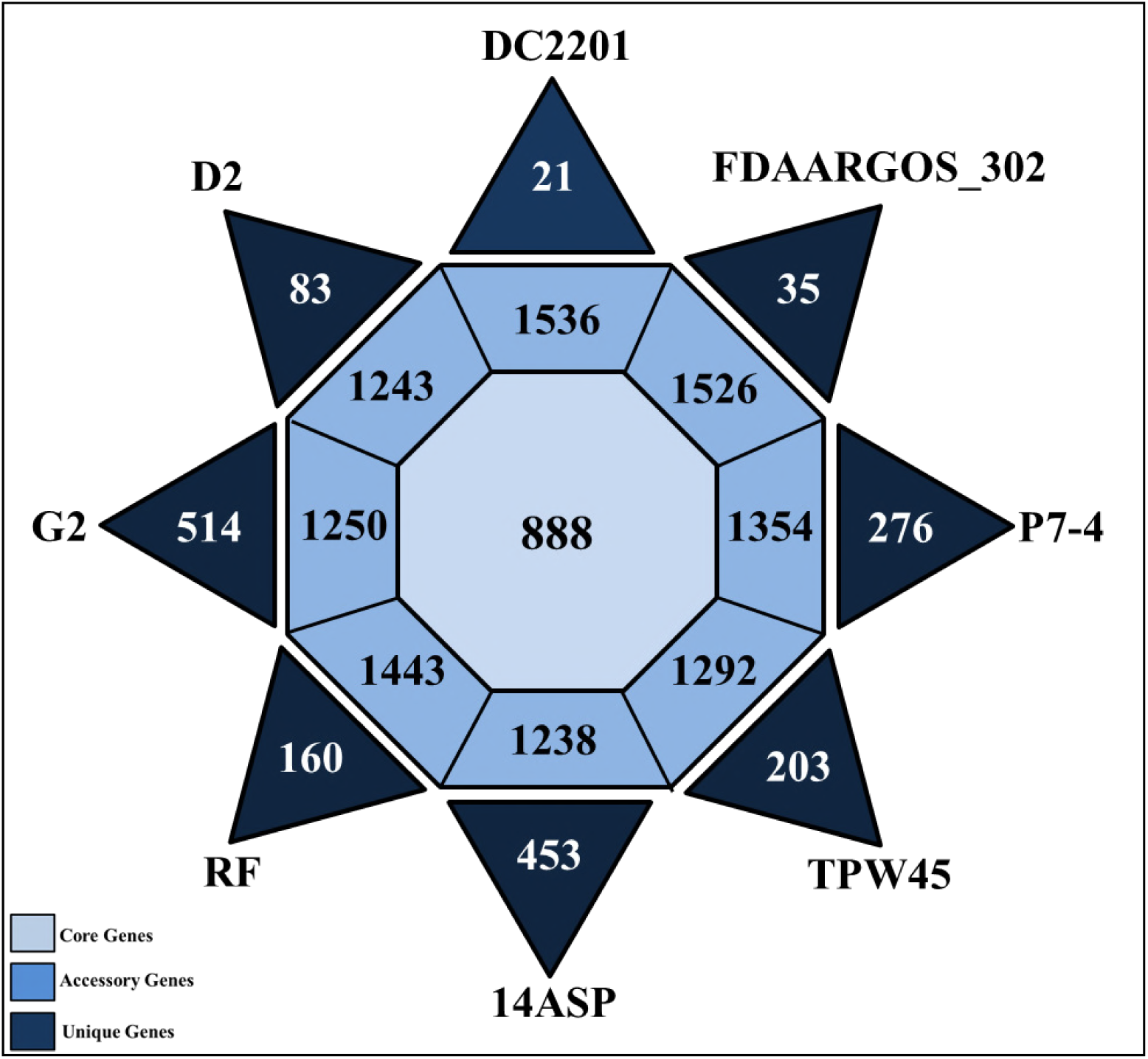
Number of Core, Accessory and Unique gene families of *K. rhizophila* genomes. The inner octagon represents the core genome consisting of 888 genes. The outer octagon represents the accessory genome and the number within them represents number of genes, while the triangles represent the unique genes associated with all the strains.

**Fig 5.**
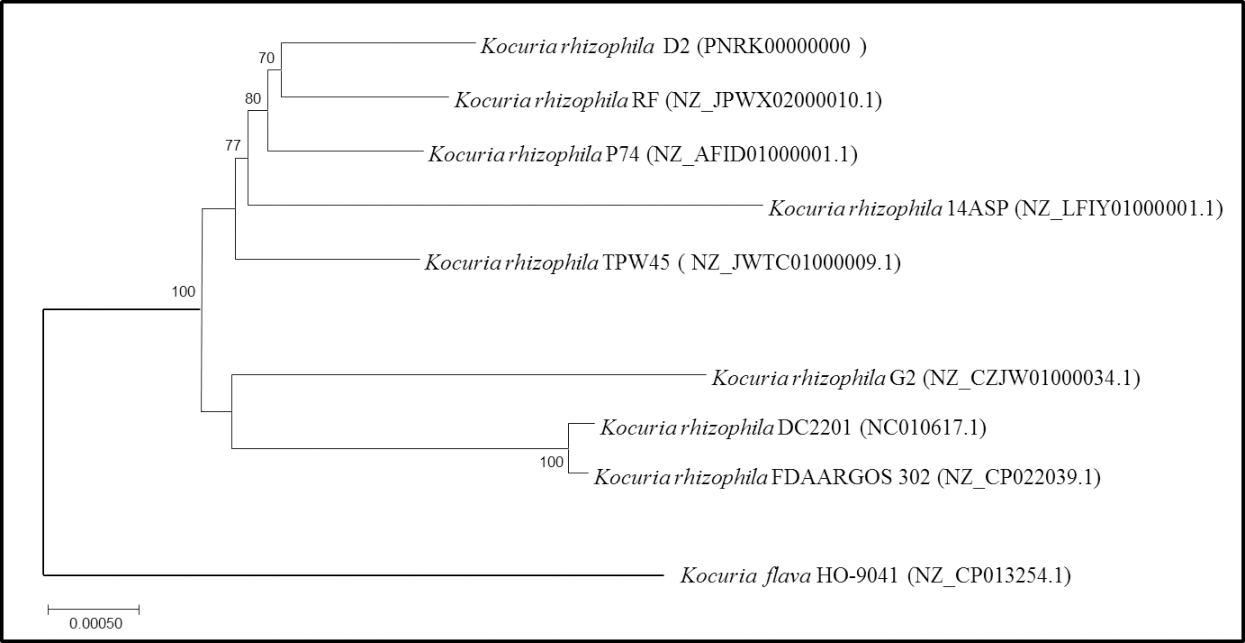
Core Genome Phylogeny. Phylogenetic tree of 8 *K. rhizophila* strains using the Maximum Likelihood method based on the GTR + G substitution model. The tree with the highest log likelihood (−17897.1414) is shown. Evolutionary analyses were conducted in MEGA6. A concatenated tree of 7909 genes was considered in the final dataset.

Functional analysis of the **accessory genes** shows the limited distribution in COG categories as opposed to core gene annotation. We could identify genes present in secondary metabolism, transport and adaption only. The functional annotation has shown the presence of a large percentage (52.19%) of genes was assigned to an uncharacterized group. The analysis also identified a number of **unique genes** associated with the various strains. The strain 14ASP had the most unique number of genes while the strain DC221 has the least number of unique genes. The annotation of these has helped to identify the role of these genes within them.

We could identify proteins involved in targeting and insertion of nascent membrane proteins into the cytoplasmic membrane, and SRP receptor for *Fts*Y and *Rec*O family involved in involved in DNA repair in strain FDAARGOS_302. In strain TPW45 arginine biosynthesis, bifunctional protein *Arg*J, O-succinylbenzoate synthase, N-acetyl-gamma-glutamyl-phosphate reductase was found. In strain D2 we could find Isopentenyl-diphosphate Delta-isomerase and general transmembrane protein while strain P7_4 had formimidoyl glutamase, single-stranded DNA-binding protein and NADH-quinone oxidoreductase subunit K. In strain 14ASP alanine racemase, cysteine--tRNA ligase, Holliday junction ATP-dependent DNA helicase *Ruv*B, crossover junction endo-de-oxyribonuclease *Ruv*C, lipoprotein signal peptidase; peptide methionine sulfoxide reductase *Msr*A and sec-independent protein translocase protein *Tat*A. While strain G2 had formamidopyrimidine-DNA glycosylase; ATP-dependent dethiobiotin synthetase *Bio*D; ribosome hibernation promoting factor; thiamine-phosphate synthase and strain RF had urease accessory protein *Ure*D while strain RF unique genes were associated with hypothetical proteins. A large portion (97.82%) of these unique genes was assigned to hypothetical proteins while only 2.17% could be assigned to some functions.

#### Mobile genetic elements (MGE)

A number of MGEs have been described in *K. rhizophila* including transposons, plasmids, and bacteriophage. Based on the screening performed the **IS element** viz. IS481, IS5, TN3 was only present in all other strains except in 14ASP (Table 5). We could identify IS21 in TPW45 and IS21 in P7-4 alone. We could identify maximum of 35 copies of intentional sequences in strain 14ASP and minimum of 7 in strain D2. Further, no **prophage** could be identified in all the eight *K. rhizophila* genomes while Clustered Regularly Interspaced Short Palindromic Repeat (**CRISPR**) were present in 3 strains: 14ASP, TPW45, RF.

**Table 5:** Distribution of IS elements within the genomes, where number indicates the copy number.

| **IS**<br>**Elements** | **DC2201** | **FDAARGOS 302** | **P7-4** | **TPW45** | **14ASP** | **RF** | **G2** | **D2** |
| --- | --- | --- | --- | --- | --- | --- | --- | --- |
| **IS1595** | 1 | 1 | 1 | 1 | 0 | 1 | 1 | 0 |
| **IS481** | 1 | 1 | 1 | 1 | 0 | 1 | 1 | 1 |
| **IS5** | 1 | 1 | 1 | 1 | 0 | 1 | 1 | 1 |
| **Tn3** | 1 | 1 | 1 | 1 | 0 | 1 | 1 | 1 |
| **IS30** | 1 | 1 | 1 | 1 | 0 | 1 | 1 | 0 |
| **ISL3** | 1 | 1 | 1 | 1 | 0 | 1 | 1 | 0 |
| **ISNCY** | 1 | 1 | 1 | 1 | 0 | 1 | 1 | 0 |
| **IS110** | 0 | 0 | 1 | 1 | 0 | 1 | 1 | 0 |
| **IS3** | 0 | 0 | 1 | 1 | 0 | 1 | 1 | 0 |
| **IS1380** | 1 | 1 | 0 | 1 | 0 | 0 | 0 | 0 |
| **IS91** | 0 | 0 | 0 | 1 | 0 | 1 | 0 | 1 |
| **IS256** | 0 | 0 | 0 | 0 | 0 | 0 | 1 | 1 |
| **IS1121** | 0 | 0 | 0 | 0 | 1 | 0 | 0 | 0 |
| **IS1649** | 0 | 0 | 0 | 0 | 1 | 0 | 0 | 0 |
| **IS1652** | 0 | 0 | 0 | 0 | 1 | 0 | 0 | 0 |
| **IS21** | 0 | 0 | 0 | 1 | 0 | 0 | 0 | 0 |
| **IS5564** | 0 | 0 | 0 | 0 | 1 | 0 | 0 | 0 |
| **IS66** | 0 | 0 | 1 | 0 | 0 | 0 | 0 | 0 |
| **ISAar35** | 0 | 0 | 0 | 0 | 1 | 0 | 0 | 0 |
| **ISAcba1** | 0 | 0 | 0 | 0 | 1 | 0 | 0 | 0 |
| **ISArsp1** | 0 | 0 | 0 | 0 | 1 | 0 | 0 | 0 |
| **ISArsp6** | 0 | 0 | 0 | 0 | 1 | 0 | 0 | 0 |
| **ISAzsp1** | 0 | 0 | 0 | 0 | 1 | 0 | 0 | 0 |
| **ISBli29** | 0 | 0 | 0 | 0 | 1 | 0 | 0 | 0 |
| **ISCgl1** | 0 | 0 | 0 | 0 | 1 | 0 | 0 | 0 |
| **ISCmi2** | 0 | 0 | 0 | 0 | 1 | 0 | 0 | 0 |
| **ISGdi10** | 0 | 0 | 0 | 0 | 1 | 0 | 0 | 0 |
| **ISJs1** | 0 | 0 | 0 | 0 | 1 | 0 | 0 | 0 |
| **ISKpn27** | 0 | 0 | 0 | 0 | 1 | 0 | 0 | 0 |
| **ISKrh1** | 0 | 0 | 0 | 0 | 1 | 0 | 0 | 0 |
| **ISKrh2** | 0 | 0 | 0 | 0 | 1 | 0 | 0 | 0 |
| **ISKrh3** | 0 | 0 | 0 | 0 | 1 | 0 | 0 | 0 |
| **ISLxc2** | 0 | 0 | 0 | 0 | 1 | 0 | 0 | 0 |
| **ISLxx4** | 0 | 0 | 0 | 0 | 1 | 0 | 0 | 0 |
| **ISMav3** | 0 | 0 | 0 | 0 | 1 | 0 | 0 | 0 |
| **ISMav4** | 0 | 0 | 0 | 0 | 1 | 0 | 0 | 0 |
| **ISMsm9** | 0 | 0 | 0 | 0 | 1 | 0 | 0 | 0 |
| **ISMysp4** | 0 | 0 | 0 | 0 | 1 | 0 | 0 | 0 |
| **ISMysp5** | 0 | 0 | 0 | 0 | 1 | 0 | 0 | 0 |
| **ISNfa1** | 0 | 0 | 0 | 0 | 1 | 0 | 0 | 0 |
| **ISPfr15** | 0 | 0 | 0 | 0 | 1 | 0 | 0 | 0 |
| **ISPfr16** | 0 | 0 | 0 | 0 | 1 | 0 | 0 | 0 |
| **ISPfr17** | 0 | 0 | 0 | 0 | 1 | 0 | 0 | 0 |
| **ISPfr19** | 0 | 0 | 0 | 0 | 1 | 0 | 0 | 0 |
| **ISPfr21** | 0 | 0 | 0 | 0 | 1 | 0 | 0 | 0 |
| **ISRae1** | 0 | 0 | 0 | 0 | 1 | 0 | 0 | 0 |
| **ISRhosp6** | 0 | 0 | 0 | 0 | 1 | 0 | 0 | 0 |
| **ISSco1** | 0 | 0 | 0 | 0 | 1 | 0 | 0 | 0 |
| **ISStpr1** | 0 | 0 | 0 | 0 | 1 | 0 | 0 | 0 |

**Genomic Islands** are distinct DNA fragments associated with mobility and we could identify maximum of 22 GIs in G2. These genomic islands comprised of a minimum of 3% to a maximum of 12.8% of genome size in the strain considered under study. These strain (DC220, FDAARGOS 302, P7-4) had an equal number of genomic islands i.e. 11. The strain D2 has the least number of islands and comprises 3.4% of the total genome. We could identify many important genes associated with a cellular function in these regions. Some of these in strain DC220 are ethanolamine permease and glutamine synthetase; dethiobiotin synthetase and fusaric acid resistance protein in strain FDAARGOS 302; acyl-CoA synthesis genes within P7-4; bleomycin resistance family protein in TPW45; glutaminase synthesis gene in 14ASP; thioredoxin in RF; short-chain dehydrogenase in G2; alkylmercury lyase (EC 4.99.1.2) in D2. Further, comparison of the genomic islands, we could not identify any common GI but could find all the IS elements, CRISPR cas genes within these regions. The genome ATLAS plot shows these differences between the strains (Fig 6).

**Fig 6.**
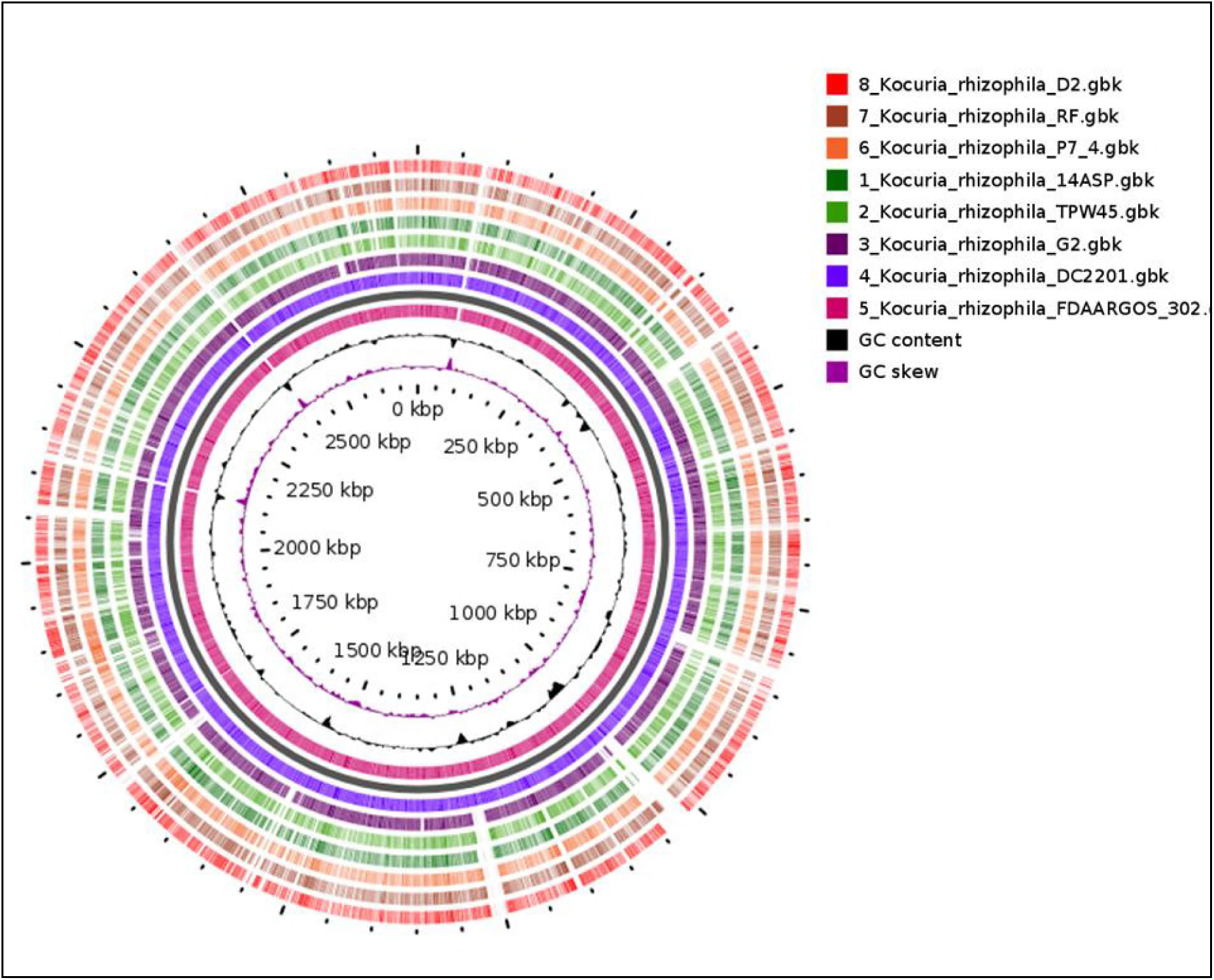
Blast Atlas of *Kocuria* genomes, with strain FDAARGOS_302 as a reference genome followed by DC2201, G2, TPW45, P7_4, RF and the outermost as D2. The difference between these genomes can be seen by the gaps in the rings.

#### Antibiotic resistance, Virulence determinants and Survival in GIT

*K. rhizophila*’s report on resistance to antibiotics is very little known. Thus in this study, we screened genomes of *K. rhizophila* against Comprehensive Antibiotic Resistance Database (CARD) for antibiotic resistance genes. Only strain RF had a single copy of beta-lactamase gene and no antibiotic resistance from other seven isolates were identified. No virulence factors could be found in any of the genomes. We could also identify genes *lytr, rrp1* for acid resistance, *clp* for bile resistance and *copA* gene for competitiveness have been identified; thus helping it to survive in the gastrointestinal tract within strain P7-4 and D2 only.

## DISCUSSION

The isolate *K. rhizophila* D2 was comprehensively characterized for its ability to colonize in the gut and, if present antibiotic resistance and virulence factors. In, the present study we use a combination of *in-vitro* and *in-silico* approaches to identify this potential of bacterial strain D2. A successful colonization in GIT by bacteria can happen if they have the ability to tolerate low pH, bile salts, oxidative stress and moreover survive in the obligate anaerobic environment (21, 22).

Principally, we tested the acid and bile tolerance and found that strain D2 was able to tolerate a low pH of 2.5 and 0.4% (w/v) bile salts with 60% and 58.8% survivability. Along with experimental evidence, the genes were identified involved in acid tolerance (*lyt, clp*) within the genome (23). We also identified genes encoding for entire primary bile salts production pathway responsible for bile salts hydrolysis activity and in-turn the tolerance. Thus indicating important characters for a bacterium to stay alive and become part of the natural GIT microbial community.

Next, we examined the aggregation and adhesion properties of strain that play an important attribute for long-term colonization in the human GIT (24). The strain D2 showed low autoaggregation capacity and adhesion. The isolate was also to adhere to human HT-29 cells which were evident from the cells surface hydrophobicity. Also, the genome of strain D2 has *copA* gene which helps in competitiveness with other bacteria (23). The resistance to hydrogen peroxide is imparted by iron-binding ferritin-like antioxidant protein and superoxide dismutase (EC 1.15.1.1) (25) found in the genome, this activity was also shown in the experiments. Overall these ability suggest its potential to thrive in the GIT conditions. Further, we could not find genes producing exopolysaccharides (EPS) and *in-vitro* results for production of EPS by strain D2 confirm its inability to produce EPS. Generally these EPS are meant to provide additional benefit as a means of protection from various stress conditions present in the human GIT (26, 27).

Genes for carbohydrate (CHO) metabolism revealed a diverse range of the gene encoding for utilizing numerous carbon sources that can be used for energy and growth. We found important genes found in CHO metabolism along with their respective transport systems. Further, the API stress assay reflected the functionally of the genome. The integrated approach of genomic and functional features of the strain together provides a comprehensive mechanism of carbohydrate utilization. The *in-silico* analysis also revealed the presence of ‘Opp proteins’ proteolytic system that help in breaking of high molecular weight proteins into smaller proteins, thus converting to absorbable forms for the human body (28–30). The utilization of gluten as a sole source of nitrogen was seen from the aminopeptidases genes present in the genome, which is a unique property for the strain D2.

The capacity to absorb sulphur, phosphate and nitrogen was found to be related with the genome by strain D2, helping them to carry out its own cellular process. The strain D2 has potential to synthesize menaquinone (vitamin K2), phylloquinone (vitamin K1), thiamine (B1) riboflavin (B2), folate (B9). These vitamins are need to be supplied exogenously and cannot be synthesized by the human cells thus essential (31, 32). Moreover, it can produce in eleven amino acids which serve as precursors for the synthesis of short-chain fatty acids (33, 34). The genome has genes for acetyl-, butyryl- and proponyl-CoA dehydrogenase but the last enzymes that convert these into acetate, buterate and propionate were absent. This is evident as this bacterium belongs to actinomycetes and acetyl-, butyryl- and proponyl-CoA dehydrogenase are further taken for another process (35, 36).

One concern regarding the strains of *K. rhizophila* is that this species has been known for its pathogenic infections. Therefore, it is of high importance to test for pathogenicity. Strain D2 showed non hemolytic activity and was sensitive to human serum and showed marginally sensitivity to some other antibiotic such as ciprofloxacin, levofloxacin, and gentamicin Thus, the strain D2 is a non-hemolytic, sensitive to major antibiotics tested and as well we could not identify the hemolytic genes in genome analysis.

Genome comparison did not reveal any significant differences (P ≤ 0.05) between the strains with reference to their GC content, genome size, an average number of genes and coding DNA sequence (CDS). The pan-genome size grew steadily with the addition of strains and the core genome stabilized, thus indicating an open pan-genome for the strains under study (Fig 4). The pan-genome analysis revealed 888 (6.57%) as core genes, 10882 (80.51%) accessory and 1745 (12.91%) as unique genes. The less number of genes in core, unique category and a large number of genes in accessory suggests the genomic fluidity of the genomes (37, 38). Further, the phylogenetic tree based on the core genome SNP based phylogeny separated 8 strains in 2 distinct clusters (bootstrap >70) (Fig 5) and no clustering based on the source of isolation.

Most of the strains under study did not harbour any antibiotic resistance gene except strain RF possessing beta-lactamase, indicating its multi-drug resistance to antibiotics such as penicillins, cephalosporins, cephamycins, and carbapenems (39, 40). No virulence genes could be identified in any strains and genes responsible for survival in GIT can be only found in strains P7-4 and D2 as these are the only gut isolates.

Insertion sequences (ISs) and bacteriophages contribute actively to bacterial evolution by integrating and exciting from the genome. In certain conditions, they provide new genetic properties such as virulence factors and antibiotic resistance (41–45). In *K. rhizophila* no such observations for IS elements with respect to virulence and antibiotic determinates could be done, also the bacteriophage did not harbour important genes related to the bacterial cellular functions. Genomic Islands are DNA fragments which usually are associated with mobility and differ between closely related strains (46, 47). These genomic islands compromised a minimum 3.2% to a maximum of 12% of the genomes in *Kocuria rhizophila* and most of these genes were assigned to hypothetical proteins. Clustered Regularly Interspaced Short Palindromic Repeat (**CRISPR**) was present in three environment strains: TPW45, 14ASP and RF. This has been attributed to the higher frequency of phage attacks present in the environments (48, 49).

In conclusion, the trio approach of *in-vitro* characterization, genome mining and comparative genomics of strain D2 have helped in the perceptive knowledge of genes responsible for surviving in the gut. Moreover, the pan-genome analysis has shown the niche-specific genes responsible for adaption and the pan-genome is open constructed on the bases of eight genomes. The unique genes present the strains D2 ability to stay within the gut and might have the potential to act as potential probiotic, as the strain produces various essential amino acids and vitamins benefiting humans. The important factor of these sequenced genomes implies the absence of virulence factors and biofilm formation ability and antibiotic resistance genes (except strain RF).

## MATERIALS AND METHODS

### Isolation and Preservation

The approval from Institutional Ethics Committee (IEC) was obtained. Three self-declared healthy volunteers were selected with consent prior to collection of the samples. We immediately transported the collected faecal samples to the lab at 4°C and processed for isolation of faecal bacteria within 6 h. Faecal samples (1g) were transferred to 9 ml of sterile saline (0.85% sodium chloride, Sigma) and mixed well (50). The serial dilutions were subsequently prepared in sterile saline, and appropriate dilutions of the samples plated on Nutrient Agar (HiMedia, Mumbai, India). Plates were incubated at 37°C for 48 h under aerobic condition. Glycerol [20% (v/v)] stocks were prepared to preserve the isolated pure cultures and froze at −80°C (50).

### Identification

Genomic DNA of the pure culture was extracted and quantified by using Qiagen Blood & Cell Culture (Qiagen, USA) and Nanodrop ND1000 (Thermo Scientific, USA) respectively. We amplified 16S rRNA gene by using universal primers, 27F (5′-AGA GTT TGA TCM TGG CTC AG-3′) and 1492R (5′-ACG GCT ACC TTG TTA CGA CTT-3′) as described in the earlier study (50). The amplified PCR products were purified using polyethylene glycol (PEG)–NaCl precipitation (50) followed by sequencing in ABI 3730xl DNA analyzer, with the help Big Dye terminator kit (Applied Biosystems, Inc., Foster City, CA). The sequence obtained was assembled using DNASTARPro, version 10 and taxonomic identity were checked by using the EZ-Biocloud server (50, 51).

### Characterization of K. rhizophila **D2**

We used Dodeca Universal I & II kit (Hi-media, India) disc diffusion method, to check antibiotic susceptibility. We tested the isolate for nine exoenzymes by standard microbiological methods viz. Phosphatase by Pikovasky’s agar base (52), lipase by tributyrin agar base (53), urease by urease agar base (54), protease by skimmed milk agar (55), gelatinase by gelatin medium (56), cellulase by CMC agar (57), amylase by starch iodine test (58), Nitrate reductase by colour change method (59) and catalase by effervescence of 6% H_2_O_2_(60). Carbohydrate utilization was conducted by HiCarbohydrateTM Kit (Sigma, India) consisting of thirty-four carbohydrates with respect to manufactures instructions. We also checked for the presence of any plasmid by Qiagen Plasmid Mini Kit (Qiagen, USA). The bile and acid tolerance assay (61–64) autoaggregation(65), cell surface hydrophobicity (66), adhesion to human HT-29 cell line (67, 68),bile-salt hydrolytic (BSH) activity (69), resistance to hydrogen peroxide (70–72) exopolysaccharide production (73), hemolytic activity (74–76), resistance to simulated GIT conditions (77–79), serum resistance (80), resistance to Simulated GI Conditions (77, 81, 82) and gluten degradation (83) was carried out as stated.

#### Statistical analysis

All the experiments were done in triplicates and the mean values and standard deviation was obtained and Duncan’s Multiple Range Test (SPSS Ver. 10.0) was used for comparisons.

#### Genome Sequencing and Assembly

We extracted genomic DNA as per the manufacturer’s protocol (QIAamp genomic DNA kit, Germany). The high-quality DNA was sequenced using Illumina MiSeq platform (2 × 300 paired- end libraries). PATRIC was used for de-novo assembly of quality-filtered reads (84, 85).

#### Bioinformatics Analyses

The draft genome sequence was annotated using RAST and the NCBI Prokaryotic Genome Annotation Pipeline (PGAP) (86). Protein coding genes, tRNA and rRNA genes from the genomes were predicted using Glimmer version 3.02 (87), tRNA_scan-SE (88) and RNAmmer (89) respectively. We also use COG database to analyze protein-coding genes (90) and Pfam domains were predicted using NCBI Batch CD-Search Tool (91). Presence of CRISPR repeats was predicted using the CRISPRFinder tool (92). We obtained open reading frames (ORFs) by using the ORF finder tool (https://www.ncbi.nlm.nih.gov/orffinder/). Prophage sequences were predicted and annotated using PHASTER (93). Bacterial insertion elements (ISs) were identified by ISfinder(94). Horizontal gene transfer was detected by genomic island tool: Islandviewer(95). Gene clusters of any bioactive compounds were identified by antiSMASH: antibiotics and Secondary Metabolite Analysis Shell (96). PlasmidFinder was used to search for plasmids within the genome (97). We used PATRIC to predict the metabolic pathways from the genome (98). Comparative genome analysis of ten whole genome sequences of *kocuria rhizophila* was done by an ultra-fast bacterial pan-genome analysis pipeline (BPGA) (99). A blast atlas was generated with the help of GVIEW Server (https://server.gview.ca/) (100).

#### Accession number(s)

We have deposited this whole genome shotgun project at GenBank under the accession PNRK00000000. The version described in this paper is version PNRK00000000.GenBank accession number for the partial 16S rRNA nucleotide sequence is MH005095.

#### Funding

This work was supported by the Department of Biotechnology, Government of India, under Grant National Centre for Microbial Resources (NCMR) (BT/Coord.II/01/03/2016).

## Acknowledgements

Additional financial support was provided by the Department of Biotechnology, Basaveshwar Engineering College, Bagalkot, under TEQUIP Grant to Bharati S. Meti.

